# Structural and functional complexity of vocalizations in a cooperatively breeding passerine, Jungle Babbler

**DOI:** 10.1101/2020.04.23.056879

**Authors:** Soniya Devi Yambem, Sonam Chorol, Manjari Jain

**Author notes:** **Corresponding Author**, Address correspondence to: Manjari Jain, Department of Biological Sciences, Indian Institute of Science Education and Research, Mohali, Punjab 140306, India.

## Abstract

Animal vocal communication ranges from simple to complex based on repertoire size, structure, and composition of calls and the information encoded in them. According to the social complexity hypothesis, communication complexity tends to increase with an increase in social complexity. While several studies on mammalian systems exist supporting this, evidence from avian systems is comparatively limited. Towards this, we present evidence for complex acoustic communication in a cooperatively breeding passerine, Jungle Babbler, based on three aspects of complexity: an extensive repertoire of acoustically-distinct calls, within-call structural complexity and the diverse behavioural contexts in which these calls are used. Jungle Babblers were found to possess a structurally and functionally diverse vocal repertoire comprising 15 different calls. Detailed acoustic analyses of multisyllabic calls revealed that these calls are composed of different notes. Further, despite a large number of notes present in the repertoire, the number of calls were limited to 15. This implies that there may be underlying rules that determine call composition to give rise to functional calls to which receivers respond. We also found that these calls were produced in a variety of affiliative and agonistic contexts and were employed towards coordination of diverse social behaviours including group movement, foraging, brood care, aggression and vigilance. Yet, 7 out of 15 vocalizations were produced in the context of vigilance. This disproportionate investment of vocalizations towards co-ordinated acoustic vigilance is characteristic of many cooperatively breeding birds. Our study extends support for the social complexity hypothesis and also lays the foundation for future investigations on combinatorial and syntactical rules underlying call structure and function in bird vocalizations.

**Significance statement:** Studies on vocal complexity in birds have focussed mainly on repertoire size, structure and function. However, fine temporal and spectral features of elements that constitute a call/song are rarely examined to evaluate vocal complexity. We examined complex communication in a cooperatively breeding social passerine, Jungle Babbler for which we assessed repertoire size, function, acoustic features of calls and of their constituent elements. Jungle Babblers were found to possess a structurally and functionally diverse vocal repertoire comprising of 15 calls, 46% of which were in the context of vigilance, thereby extending support to the social complexity hypothesis. We also found that several calls were composed of multiple, acoustically distinct notes. These findings will be foundational in understanding the interrelations between sociality and communicative complexity and underlying combinatorial rules that determine call structure and function.

## Introduction

Theories on the evolution of language have long acknowledged the importance of sociality in the evolution of complex communication. The social complexity hypothesis posits that communication complexity tends to increase with an increase in social complexity (Blumstein and Armitage 1997). This is because social animals are likely to require higher sophistication in communication to coordinate different behaviours and sustain the relationships between individuals within a social group. The complexity of communication is determined by both structural and functional complexity. Structural complexity of vocalizations is determined by the compositional features of calls (how many elements make a call and how these elements are organized within a call), overall acoustic features of the call (temporal and spectral characteristics) and the total number of unique vocalizations of a species (Hailman and Ficken 1986; Zimmermann and Lerch 1993; reviewed in Freeberg et al. 2012). Additionally, the different behavioural contexts in which calls differ also add to the communication complexity (reviewed in Freeberg et al. 2012; Crane et al. 2016). Based on the number of elements in a call, avian vocalizations can be classified as monosyllabic or multisyllabic (Rothstein 1988). A ‘syllable’ or ‘note’ is the simplest element of a call and forms the smallest acoustic unit, which when combines with other such units (of the same or different type) forms a ‘phrase’ (multisyllabic call). The total vocalization produced by an animal throughout its lifetime is called a vocal repertoire (Searcy 1992). Repertoire size vary from species to species, for instance, male Common Grackles have only one song type (Searcy 1992) whereas Common Nightingale sings up to 180 different song types (Weiss et al. 2014). A smaller repertoire is considered as less complex compared to a larger repertoire size (Blumstein and Armitage 1997).

The function of the call repertoire is different in different species and requires extensive behavioural observations to ascertain. In many species of songbirds, complex vocalizations serve as a sexual display to attract mates or as territorial display against conspecific and heterospecific individuals. It has been shown that individuals with larger repertoire sizes are more successful in attaining mates (Catchpole 1987; Robinson and Creanza 2019) or can hold a territory for a longer duration (Hiebert et al. 1989). It has also been reported that in several species the size of the repertoire increases with complexity in vocalization context. For instance, Diana monkeys produce predator-specific alarm calls (Zuberbühler 2002) whereas, in Meerkats, alarm calls vary depending not only on predator type but also on the level of urgency (Manser 2001).

Social animals exhibit higher frequency and diversity of interactions between different individuals than solitary animals and possess a larger and more complex repertoire to deliver meaningful inference (Freeberg et al. 2012; Krams et al. 2012; Leighton 2017). Based on the interaction type, calls are broadly classified as either ‘affiliative’ or ‘agonistic’. Calls that are used in maintaining social bonds are ‘affiliative’ whereas ‘agonistic calls are produced against conspecific and heterospecific rivals (Kondo and Watanabe 2009). Quantifying and categorizing the entire repertoire based on signal structure and function can provide valuable insights into the behaviour of the species and selective forces that drive the evolution of complex communication.

Jungle Babblers (*Turdoides striata*) are cooperatively breeding passerines that live in groups of 3-20 individuals and are widely distributed throughout lowland India, both in rural and urban habitats (Andrews and Naik 1970; Gaston 1977; Ali and Ripley 1978). Group members engage in many social behaviours such as cooperative brood care, sentinel duty, collective foraging, anti-predator behaviour, allogrooming and intergroup confrontations. (Andrews and Naik 1970; Gaston 1977). To maintain bonding within the social group, it is expected that there would be vocalizations associated with interactive behaviours, thereby raising the possibility that Jungle Babblers possess a complex communication system. Andrew and Naik (1970) and later Gaston (1977) provided onomatopoeic descriptions of some vocalizations of the species and the situation in which the calls were observed, thereby raising the possibility of complex acoustic communication in this cooperative breeder. However, so far, no study has examined complex acoustic communication in Jungle Babblers, despite their broad distribution, the potential for multiple vocalizations and known social system. In this study, we aim to examine complex acoustic communication in Jungle Babblers by a quantitative and systematic investigation of the vocal repertoire of the species. The major objectives were 1. To quantify the acoustic structure of calls to estimate the number of structurally distinct call types, 2. To determine the acoustic features of the constituent elements (notes / syllables) of each of these call types 3. To ascertain the behavioural context associated with each call type through extensive behavioural observations. This study provides the first acoustic characterization of the vocalization of Jungle Babblers, carried out both across and within call type, and the associated behavioural contexts. Apart from providing the bedrock for future investigations of vocal complexity in this species, this study will allow a comparative investigation of complex communication in avian systems with varying degrees of sociality and ecological conditions.

## Materials and methods

Since the data collected for this study was completely based on observations and recordings of animals in the field, it was not possible to record data blind.

### Study site

The study was conducted in Mohali region, located in the eastern part of Punjab in northern India (30°36’ and 30°45’N latitude and 76°38’and 76°46’E longitude), which covers an area of about 116.50 km^2^. According to the Koppen-Geiger climate classification system, the climate of Mohali comes under the ‘Cwa’ category (Kottek et al. 2006). Mohali has a humid subtropical climate that is variable throughout the year with a hot summer and cold, dry winter separated by a brief period of tropical monsoon climate. The habitat of the study site comprised of shaded gardens and closed-canopy woodland with trees such as *Populous deltiodes, Ficus religiosa, F. glomerata, Vachellia nilotica, Morus alba, M. nigra*, and *Psidium guajava, Leucaena leucocephala, Chukrasia tabularis, Callistemon sp*., and shrubby habitats dominated by bushes of *Lantana camara*, *Ricinus communis* and *Cannabis sp*. (Fig. S1).

### Behavioural data collection

The fieldwork for this study was conducted between May 2016 and March 2020, during which we carried out systematic behavioural observations to interpret the context of vocalizations of Jungle Babblers. All behavioural observations were made on free-ranging birds using 8 x 42 binoculars (Nikon, Monarch 7) following *Ad libitum* and focal animal sampling (Altman 1974). Towards this, the vocalizing individual was identified and its behaviour was noted. This was accompanied by noting the behaviour of all group members in sight and a quick scan of the environment. Any response given to the signaller was also noted.

### Acoustic data collection

Recordings were taken from free-ranging Jungle Babblers in their natural habitat at a distance of up to 10 meters from the caller. A solid-state recorder (Marantz PMD661 *MKII*; frequency response: 20 Hz - 20 kHz), connected to a super-cardioid shotgun microphone (Sennheiser ME66 with K6 PM; frequency response: 40 Hz to 20 kHz), covered with a foam windscreen (Sennheiser MZW66) was used to record all vocalizations (sampling rate of 44.1KHz and 16-bit accuracy). Calls of nestling were recorded while they were inside the nest and from fledgling outside the nest. Fledglings were identified based on iris colour (juvenile - black; adult - pale white; Fig. S2). Recordings were also made from individuals trapped in mist-net and during their subsequent release. To minimize the overlap of calls between individuals, recordings were focused on a single individual except for chorus calls. While recording the vocalizations, the behaviour of the caller and receivers were noted as described above and the surroundings were scanned. These observations were announced at the end of the call recording. Data from the behavioural observations were then used to interpret the context of vocalizations and the recordings were categorized under these behavioural categories. Andrew and Naik (1970) and Gaston (1977) catalogued a list of situations in which Jungle Babblers vocalize. This list served as a valuable reference library for validating our independent inferences.

### Acoustic analyses

A total of 303 recordings comprising of 1895 calls were processed in Raven Pro 1.5 (Cornell Laboratory of Ornithology, USA) and spectrograms were generated using Hann window function, size 512 with a 50% overlap. After generating the spectrograms, only those calls with no or minimum overlap were retained for further analysis except in the case of chorus calls. For the acoustic categorization of calls, as a first step, calls were classified under the same or different call type based on aural and visual inspection of the spectrograms. Calls were also examined based on the inter-note interval. If the time gap between two notes was <= 0.1 s, they were considered to belong to the same call (Catchpole and Slater 2010). This allowed us to categorize all call types under one of two categories: ‘monosyllabic’ (single note call) or ‘multisyllabic’. The third category of calls called ‘chorus calls’ included those vocalizations in which multiple individuals vocalized simultaneously. Within each of these three categories detailed acoustic characterization of different calls was then carried out based on 7 different acoustic features: A) spectral parameters i) frequency 5% (Hz) ii) frequency 95% (Hz) iii) bandwidth 90% (Hz) and iv) peak frequency (Hz); B) temporal parameters: i) call duration (s) and ii) inter-note/syllable interval (s) iii) total number of notes in a call. Finally, all multisyllabic calls were subjected to further analyses wherein a total number of notes in each call was identified and acoustic analyses of each constituent note were carried out based on all spectral parameters and note duration. For the analyses of the chorus calls, only spectral parameters were considered as signal overlap from different callers made analyses of temporal features unreliable.

### Statistical analysis

Calls that were preliminarily categorized under different call types based on behavioural observation and audio and visual observation of the spectrogram. They were then subjected to rigorous statistical analyses based on their acoustic features, as determined by the signal analyses carried out. For this, pairwise comparisons were carried out between calls belonging to the same category (monosyllabic, multisyllabic or chorus calls). Calls were considered to be different from each other if they differed significantly in at least 1 out of the 7 acoustic parameters examined. Further, within call complexity was examined for all multisyllabic calls by examining whether a given call was composed of multiple, acoustically distinct notes or was composed of repeats of the same elements. All the statistical analysis was carried out in Statistica 64 (Dell Inc.2015, Version 12). To check for normality of data, the Shapiro-Wilk’s test was used. ANOVA, and Kruskal-Wallis ANOVA were performed for data that followed and did not follow a normal distribution, respectively. Further, for pairwise comparisons, unpaired t-test for normally distributed data and Mann-Whitney U (MW U) test for data that did not follow normal distribution were carried out. This allowed us to examine if the behavioural categorization of calls as different call types was consistent with the acoustic characterization and to examine the complexity of vocal repertoire both across and within call types.

## Results

### Vocal repertoire and structural complexity of calls

The audio-visual inspection of the spectrogram and analyses of inter-note interval resulted in Jungle Babbler vocalizations being classified into 15 different call types. Adult vocalizations included 5 monosyllabic, 6 multisyllabic and 2 chorus calls, whereas juvenile vocalizations included 2 call types, both of which were monosyllabic (Fig. 1, Table 1). All monosyllabic adult calls were found to be significantly different from each other based on their acoustic features (Kruskal-Wallis ANOVA; Table S1a). The pairwise comparison revealed that all calls were significantly different from each other by at least three acoustic parameters (MW U test; Table S1b, Fig. 2a-e). The two Juvenile calls were significantly different from each other only in temporal characteristics and not based on any spectral parameters (t-test and MW U test; Table S2, Fig. 2f).

**Fig.1.**
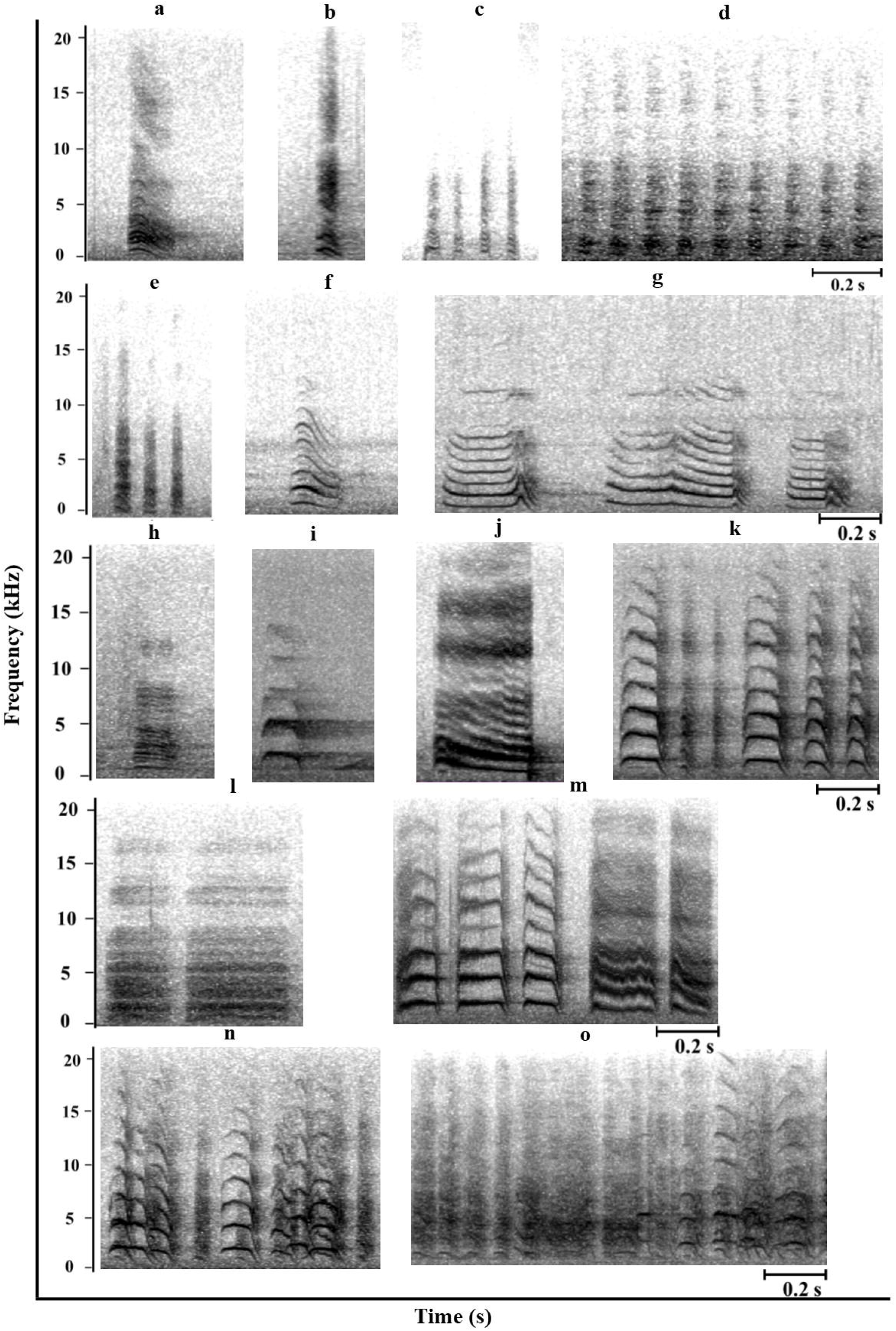
Spectrograms of different call types of Jungle Babbler. **Affiliative calls: a** contact; **b** foraging; **c** prompt; **d** prompt flight; **e** flight; **f** fledgling close and **g** begging. **Agonistic calls: h** sentinel soft; **i** threat; **j** distress; **k** alert; **l** harsh; **m** intermediate alert; **n** mobbing and **o** intergroup fight

**Fig. 2.**
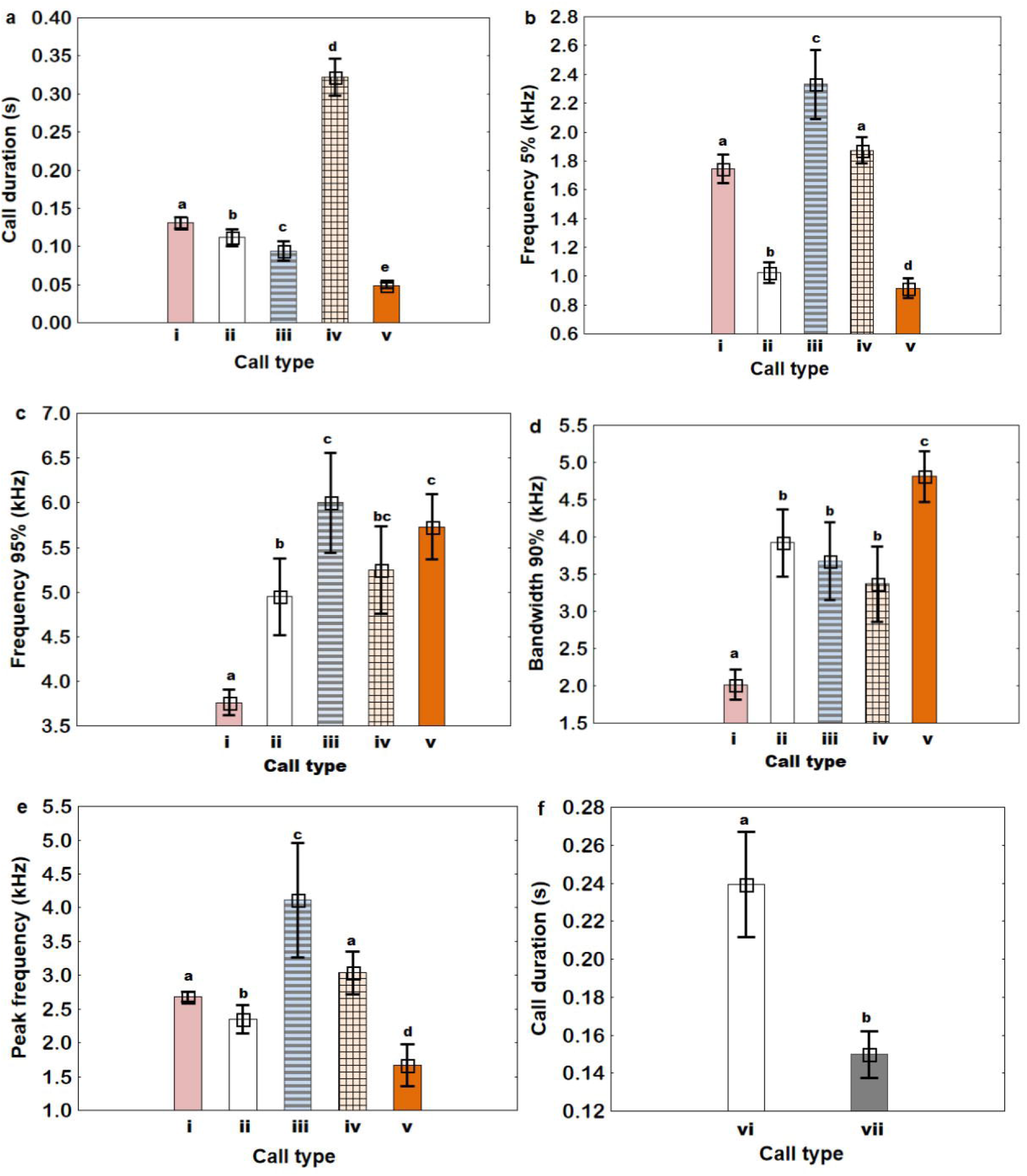
Variation in different monosyllabic call types of adult and juvenile of Jungle Babblers based on 5 different acoustic parameters. **Adult calls: a** call duration; **b** frequency 5%; **c** frequency 95%; **d** bandwidth 90% and **e** peak frequency. **Juvenile calls: f** call duration. Roman numbers on the x-axis correspond to different call types: **i** contact; **ii** sentinel soft; **iii** threat; **iv** distress; **v** foraging; **vi** begging; and **vii** fledgling close. Different alphabets represent the statistical difference between call types. Bar corresponds to mean with 95% confidence interval

**Table 1.** Temporal and spectral characteristics of 15 vocalizations type of Jungle Babbler showing mean ± std. N and n correspond to the number of recordings and calls for each call types respectively. Blue colour filled cells represent affiliative call category and unfilled represent agonistic call category. * represent onomatopoeic description used by Gaston (1977)

| S. no | Call types |  | N | n | No. of notes | Call duration (s) | Inter-note interval (s) | Frequency 5% (kHz) | Frequency 95% (kHz) | Frequency bandwidth 90% (kHz) | Peak frequency (kHz) |
| --- | --- | --- | --- | --- | --- | --- | --- | --- | --- | --- | --- |
|  | Behavioural context | Onomatopoeic description |  |  |  |  |  |  |  |  |  |
| 1 | Contact | Chack * | 33 | 175 | 1 | 0.13 ± 0.02 | - | 1.75 ± 0.28 | 3.76 ± 0.41 | 2.02 ± 0.55 | 2.68 ± 0.22 |
| 2 | Fledgling close | Fledgling chack | 10 | 41 | 1 | 0.15 ± 0.02 | - | 1.58 ± 0.26 | 4.28 ± 0.98 | 2.70 ± 1.01 | 2.66 ± 0.34 |
| 3 | Begging | Begging | 14 | 78 | 1 | 0.24 ± 0.05 | - | 1.42 ± 0.4 | 4.82 ± 0.83 | 3.40 ± 0.98 | 3.01 ± 1.05 |
| 4 | Foraging | Cuk * | 17 | 63 | 1 | 0.05 ± 0.01 | - | 0.92 ± 0.13 | 5.73 ± 0.71 | 4.81 ± 0.67 | 1.67 ± 0.60 |
| 5 | Prompt | Ca-ca-ca | 19 | 47 | 4.45 ± 1.4 | 0.31 ± 0.13 | 0.04 ± 0.01 | 0.91 ± 0.19 | 5.75 ± 1.40 | 4.84 ± 1.36 | 1.59 ± 0.35 |
| 6 | Flight | Cu-cu-cu * | 31 | 79 | 3.68 ± 0.65 | 0.30 ± 0.06 | 0.05 ± 0.01 | 0.89 ± 0.28 | 4.26 ± 0.67 | 3.36 ± 0.80 | 1.81 ± 0.45 |
| 7 | Prompt flight | Long cu-cu-cu | 22 | 60 | 7.07 ± 3.88 | 0.55 ± 0.31 | 0.04 ± 0.01 | 0.92 ± 0.20 | 4.60 ± 0.97 | 3.68 ± 1.06 | 1.67 ± 0.35 |
| 8 | Threat | Shriek * | 12 | 17 | 1 | 0.1 ± 0.02 | - | 2.33 ± 0.38 | 6.00 ± 0.88 | 3.67 ± 0.82 | 4.11 ± 1.33 |
| 9 | Sentinel soft | Low chack | 21 | 99 | 1 | 0.1 ± 0.02 | - | 1.02 ± 0.16 | 4.95 ± 0.95 | 3.92 ± 0.99 | 2.35 ± 0.46 |
| 10 | Distress | Kya-kya-kya | 22 | 220 | 1 | 0.32 ± 0.06 | - | 1.88 ± 0.20 | 5.25 ± 1.1 | 3.37 ± 1.14 | 3.04 ± 0.71 |
| 11 | Alert | Cackle * | 23 | 363 | 4.14 ± 2.91 | 0.54 ± 0.34 | 0.05 ± 0.01 | 1.58 ± 0.33 | 6.46 ± 0.72 | 4.87 ± 0.76 | 3.45 ± 0.95 |
| 12 | Harsh | Khack | 38 | 332 | 1.57 ± 0.78 | 0.34 ± 0.25 | 0.07 ± 0.02 | 1.36 ± 0.30 | 5.50 ± 1.17 | 4.14 ± 1.02 | 2.73 ± 0.85 |
| 13 | Intermediate alert | Khack cackle | 19 | 299 | 2.98 ± 1.26 | 0.90 ± 0.30 | 0.06 ± 0.01 | 1.67 ± 0.21 | 5.81 ± 0.79 | 4.14 ± 0.67 | 3.27 ± 0.80 |
| 14 | Mobbing | Wheezy cackle * | 15 | 15 | - | - | - | 1.85 ± 0.12 | 6.40 ± 0.55 | 4.55 ± 0.53 | 4.12 ± 0.58 |
| 15 | Inter-group fight | Guttural noise * | 7 | 7 | - | - | - | 1.63 ± 0.31 | 6.05 ± 0.38 | 4.42 ± 0.40 | 3.89 ± 0.48 |

Similarly, there was a significant difference between all multisyllabic calls based on their acoustic features (Analysis of variance; Table S3a and Kruskal-Wallis ANOVA; Table S3b; Fig. 3). The pairwise comparison showed that all multisyllabic calls are also significantly different from each other (t-test and MW U test; Table S3c and d Fig.3) by more than one parameter.

**Fig. 3.**
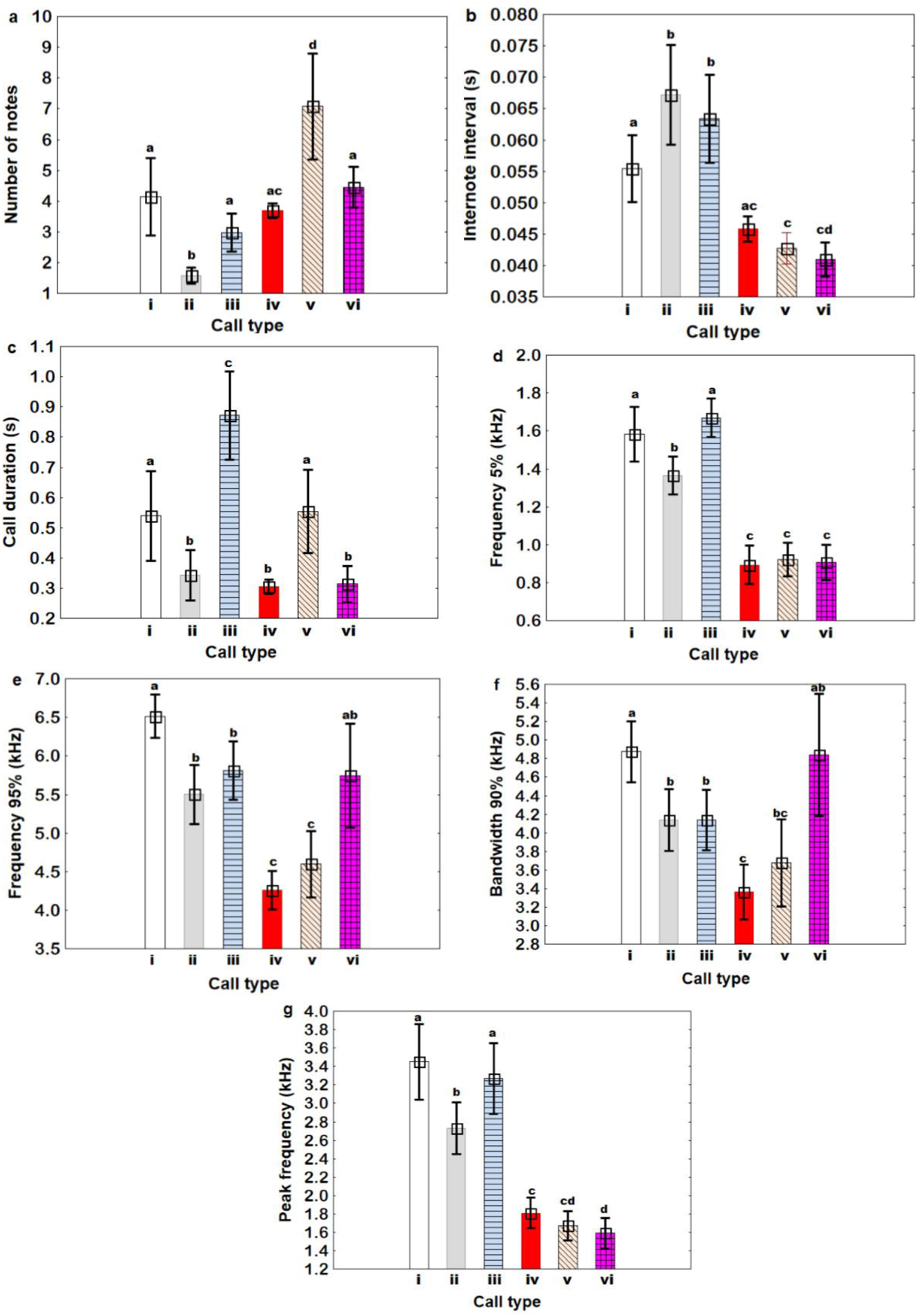
Variation in different multisyllabic calls of adult based on 7 acoustic parameters. **a** number of notes; **b** inter-note interval; **c** call duration; **d** frequency 5%; **e** frequency 95%; **f** bandwidth 90%; and **g** peak frequency. Roman numbers on the x-axis correspond to different call types; **i** alert; **ii** harsh; **iii** intermediate alert; **iv** flight; **v** prompt flight; and **vi** prompt. Different alphabets represent the statistical difference between call types. Bar corresponds to mean with 95% confidence interval

The two chorus calls were found to be not significantly different based on their spectral features (t-test and MW U test, Table S4a and b).

Based on aural and visual inspection, we found that each multisyllabic call was composed of different note types, which were then compared statistically within a call type. Multisyllabic calls were found to be composed of at least three acoustically distinct note type each (Analysis of Variance, Kruskal-Wallis ANOVA and MW U test; Table S5a-c; Fig. 4 and 5).

**Fig. 4.**
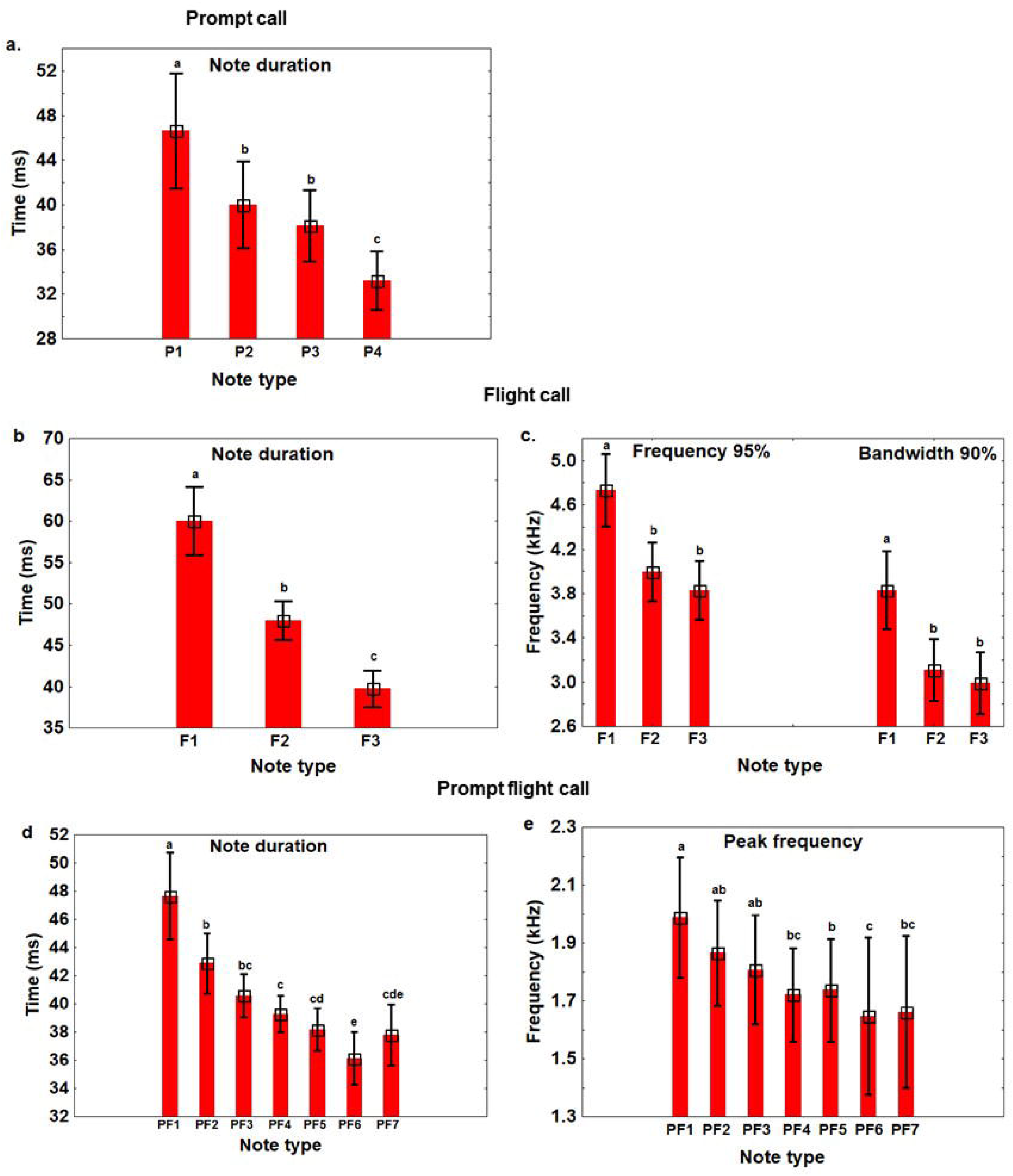
Variation in notes within multisyllabic affiliative call. **a** note duration of 4 note types of prompt call; **b** note duration and **c** frequency 95% and bandwidth 90% of 3 note types of flight call; and **d** note duration and **e** peak frequency of 7 note types of prompt flight calls. Different alphabets represent the statistical difference between call types. Bar corresponds to mean with 95% confidence interval

**Fig. 5.**
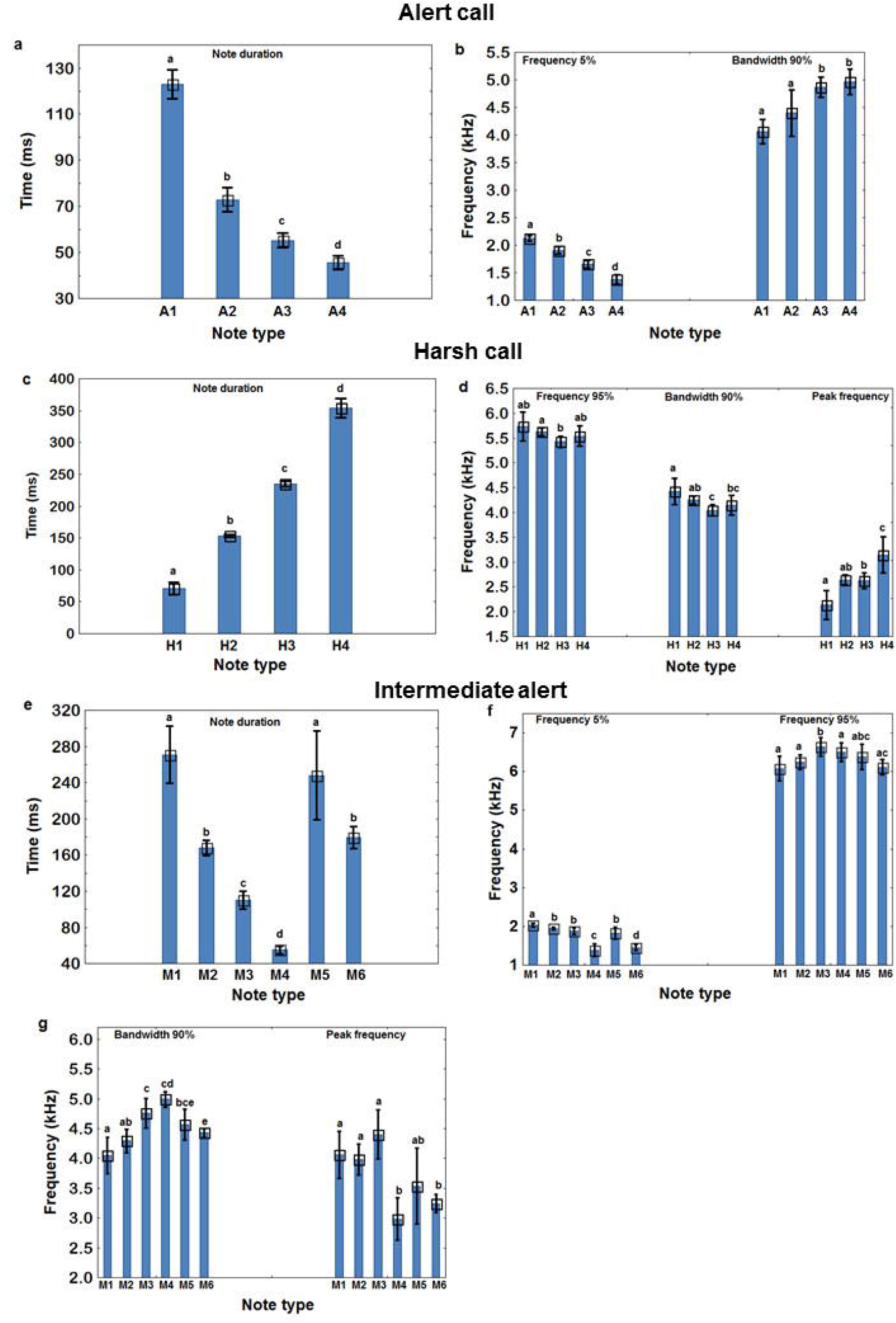
Variation in notes by temporal and spectral parameter within multisyllabic agonistic calls. **a** note duration and **b** frequency 5% and bandwidth 90% of 4 note types of alert call; **c** note duration and **d** frequency 95%, bandwidth 90% and peak frequency of 6 notes of harsh call; and **e** note duration, **f** frequency 5% and frequency 95% and **g** bandwidth 90% and peak frequency of 6 note types of intermediate alert call. Different alphabets represent the statistical difference between call types. Bar corresponds to mean with 95% confidence interval

### Behavioural context of vocalizations

This large and structurally complex vocal repertoire of Jungle Babblers was also found to be functionally diverse. The results from the behavioural observations suggest that these vocalizations are produced in 2 major functional contexts: affiliative and agonistic. Detailed below are the behavioural contexts in which the signallers were observed to produce these diverse vocalizations.

### Affiliative calls

i) Contact (Chack) call: This call is used to contact conspecifics and is produced when an individual is separated from the group. In response to this call, a member of the group gives the same call back and eventually approached the left behind signaller leading to a reunion of a diverted individual with the rest of the group. This is a loud call that consists of only one note (monosyllabic) with faintly visible harmonics (Fig. 1a).
ii) Foraging (Cuk) call: This is a very soft call produced by one or more individuals of the group while foraging on the ground/leaf litter. No unique behaviour or reaction was observed when this call is produced and the group continues to forage. This monosyllabic call is the call with the shortest duration (Fig. 1b).
iii) Prompt (Ca-ca-ca) call: This call is given by an adult heading towards nest for food provisioning. Nestlings respond to this call with a ‘begging call’ (described later). The call is multisyllabic, mainly composed of 3 - 4 notes that are similar spectrally, however, note duration decreases progressively (Fig. 1c).
iv) Prompt flight (Long cu-cu-cu) call: This vocalization is also made when adults are in proximity to fledglings, typically around nests. While producing these calls, adults take short flights from one position to another and back to the fledgling. This seems to prompt fledgling movement. Fledglings were observed to move out of the nest or move from their current location to another in response to this call by adults. This multisyllabic call is usually produced by more than one individual at a time and is composed of the most number of notes (3 - 11 notes per call). The notes at the beginning of the call are of longer duration and higher peak frequency and forthcoming notes show a gradual decrease in both these parameters (Fig. 1d).
v) Flight (Cu-cu-cu) call: This call is generally produced during group displacement. The call is initiated by one individual and is eventually joined in by the rest of the group members. Group movement initiates soon after. This is a soft multisyllabic call composed of 3 - 4 notes. Here too, notes in the beginning, have a longer duration and higher frequency 95% while trailing notes progressively become shorter in duration and have lower values for frequency 95% (Fig. 1e).
vi) Fledgling close (Chack) call: This vocalization is produced by fledglings or juveniles in close proximity to adults. Adults do not respond to this call. Spectrally, this monosyllabic call is almost similar to adult contact call however, adult contact calls are shorter in duration than these calls (Fig. 1f).
vii) Begging call: Nestlings and fledglings produce this monosyllabic call during food provisioning and in response to prompt calls. Spectrally, this call is very dynamic. This call is generally accompanied with the wide opening of beak and rigorous flapping of wings (Fig. 1g).

### Agonistic calls

viii) Sentinel soft (Low chack) call: This monosyllabic call is exclusively produced by a sentinel-an individual on vigilance duty, on an elevated perch, forgoing foraging while the rest of the group members forage, Andrews and Naik (1970). While producing this vocalization, the caller is generally not very alert and may even groom at times. There is no visible change in the behaviour of group members in response to this call (Fig. 1h).
ix) Threat (Shriek) call: This call is produced in response to the sudden approach of a potential threat. Both the caller and receiver usually respond instantly with a startle and take shelter in the closest tree or foliage. After a while, inside the cover, they might start grooming or allogrooming or resume regular behaviour (generally foraging). This call is monosyllabic but sometimes two notes may be produced in continuation. Among all monosyllabic calls, this call has the highest peak frequency (Fig. 1i).
x) Distress (Kya-kya-kya) call: This is a loud monosyllabic call produced repeatedly by an individual in distress. Such a situation may arise when an individual gets trapped in mist-net or is handled by a human or a predator. This call attracts group members towards the caller. Among all monosyllabic call, this call has the highest note duration (Fig. 1j).
xi) Alert (Cackle) call: During intrusion by any potential threat, a perched individual produces this call, accompanied by periodic hops from one side to the other, flapping of wings and twitching of tail up and down. This call may induce other members of the group to join in the vocalization, whereas on other occasions no visible response from the group members is observed. This call is a multisyllabic call comprising of 2 - 4 different notes. As the call proceeds, there is a decrease in note duration and minimum frequency (frequency 5%) whereas the frequency bandwidth (bandwidth 90%) increases progressively (Fig. 1k).
xii) Harsh (Khack) call: This is a noisy harsh call, produced by any perched individual in response to any intrusion which is not an immediate threat, including the observer. This call is also produced while individuals are foraging in the foliage or are in the queue for food provisioning. There was also no visible response by group members in response to this call. This call is multisyllabic and comprises of 1-5 notes forming a phrase (Fig. 1l).
xiii) Intermediate alert (Khack cackle) call: Any perched individual produces this call in the presence of the observer but also for other intruders. The intensity of the caller’s behaviour is somewhat intermediate to the harsh and alert call. No visible response was observed from the group. The call comprises of 6 structurally different notes/syllables with highest call duration (Figure 3j and 3k). Moreover, there is no fixed pattern in the composition of the call (Fig. 1m).
xiv) Mobbing (Wheezy cackle) call: This vocalization is a chorus call wherein more than one individual at a time is involved. It was observed to be produced in the presence of an immediate and proximate predator such as domestic/feral cats, Indian Grey Mongoose, Greater Coucal, Spotted Owlet, Barn Owl, snakes, and Bonnet Macaque. While making this call, all individuals involved, flutter around and ‘harass’ the potential predator until the predator retreats. However, if the predator did not move from its position, then the group disperses from the location. This vocalization also attracts neighboring heterospecific birds to join in mobbing the predator accompanied by loud and urgent vocalizations. It may, however, be noted that while producing mobbing calls, Jungle Babblers were not observed making any physical interactions with the predator (Fig. 1n).
xv) Intergroup fight (guttural noise) call: This is also a chorus call, produced when two different groups of Jungle Babblers come in contact with each other. They produce this vocalization while moving from one tree to another, chasing each other. Along with this vocalization, sometimes individuals engage in mid-air physical fights, striking each other using beaks and claws while falling to the ground, after which they usually disengage. Concurrently other members of the groups surround the fighting pair while producing this vocalization. They also make this call by positioning themselves facing each other on different trees without physical interaction. It may be noted, however, that intergroup fights are rare (Fig. 1o).

## Discussion

The major drivers of vocal complexity in avian systems include sexual selection and sociality (MacDougall-Shackleton 1997; Freeberg et al. 2012) and measures of complexity include both structural (repertoire size and features of calls) and functional (behavioural contexts) aspects of vocalizations (Crane et al. 2016; Holt 2017). Cooperative breeding has been found to be a strong predictor of large repertoire size in avian systems (Leighton 2017) and several avian cooperative breeders possess multiple structurally-distinct calls (Ficken et al. 1978; Seddon 2002; Warrington et al. 2014; Crane et al. 2016). Evaluation of overall structural complexity, however, must also incorporate an assessment of the fine structure of all vocalizations by measuring the temporal and spectral features of elements (notes/syllables) that constitute a call/song. This level of analysis is largely missing in the assessment of vocal complexity in social birds (Greig et al. 2008; Grieves et al. 2015; Crane et al. 2016).

In this study, we examined communicative complexity in Jungle Babblers by examining repertoire size, overall acoustic features of all calls, and their constituent elements, and the behavioural contexts in which vocalizations are produced. The vocal repertoire of Jungle Babblers comprises of 7 monosyllabic and 8 multisyllabic calls. All calls were structurally distinct from each other. Note-level analyses within multisyllabic calls revealed that each call were constructed by the combination of several acoustically distinct notes (call type (number of distinct notes): prompt (3), flight (3), prompt flight (4), alert (4), harsh (4), intermediate alert (6)). While 31 notes were recorded in the vocal repertoire of this species, the functional repertoire size is limited to 15 calls. This implies that this large variety of notes combine in a limited number of ways to form meaningful calls. It is also possible that some of the notes are shared across different calls and examining this would require across call note comparisons. This study thereby lays the foundations to further investigations on understanding the limits of combinatorial rules that determine the composition of meaningful multisyllabic calls in this social bird.

With respect to the functional aspects of the vocal repertoire, our findings are consistent with the observations of Andrews and Naik (1970) and Gaston (1977) that various affiliative and agonistic behaviours are mediated by vocalizations in Jungle Babblers. Affiliative calls include those that coordinate group movement as well as food provisioning and brood care. For instance, flight call in Jungle Babblers clearly induces all group members to cohesively displace to a new location and contact call functions to reunite lost or left-behind members with the rest of the group. Calls like prompt and prompt flight induce fledglings to beg and fly, respectively. Similar findings have been reported in Pied Babblers that produces ‘purrs’ that prompt fledglings towards a food source and ‘clucks’ to induce group displacement (Engesser et al. 2017). The exact function of the foraging call in Jungle Babblers remains unclear. We speculate that similar to foraging calls in Pied Babblers, this call may play a role in maintaining the spacing between the foragers in order to enhance foraging efficiency (Radford and Ridley 2008).

Our findings also suggest that 8 out of 15 vocalizations of Jungle Babblers are produced in agonistic context towards conspecifics and heterospecifics. It includes inter-group fight calls (guttural noise) produced against conspecific rivals and 6 others, including threat, distress, harsh, intermediate alert, alert, and mobbing calls that are produced against heterospecifics, mainly potential predators. This disproportionately high representation of calls towards vigilance could be because predation imposes a strong selective pressure by directly impacting the fitness of an individual (Leighton 2017). It is known that vigilance behaviour is costly since the animals must interrupt foraging in order to scan the environment for predators (Wickler 1985). By having coordinated acoustic vigilance, individuals can mitigate this cost as it renders visual scans by every individual unnecessary. This, however, is contingent on the animals producing reliable alarm calls, allowing receivers to choose the appropriate escape strategy, thereby increasing the chance of survival (Marler 1967). Thus, it would be useful to have functionally referential alarm vocalizations that inform the receivers about the category of predator (aerial or terrestrial or even predator specific; Seyfarth et al. 1980; Naguib et al. 1999). It would also be useful to encode the level of urgency (encoding distance of potential threat) to allow the receivers to respond in an appropriate time (Manser 2001). All of these together is likely to increase the proportional representation of vigilance vocalization in the vocal repertoire of cooperative breeders. Meerkats, for instance, are known to possess 30 different vocalizations of which 18 (60%) are produced in the context of vigilance. Similar to our findings, studies on other cooperatively breeding birds have shown that a large number of vocalizations are dedicated to vigilance. This includes Chestnut-crowned Babblers (4 out of 13 calls; Crane et al. 2016), Pale-winged Trumpeter (5 out of 12 calls; Seddon et al. 2002) and Black-capped Chickadees (5 out of 11 calls; Ficken et al.1978). In a study carried out on 253 bird species across the globe, representing 59 families, it was found that cooperative breeders dedicated a significantly higher proportion of their vocal repertoire to vigilance related vocalizations (Leighton 2017).

Alarm calls can be categorised into three main types: flee, mob and distress (reviewed in Magrath et al. 2015). Flee-type calls are generally short duration pure tone calls making them difficult to localize (Marler 1955). In Arabian Babblers, alarm calls produced by foragers were found to be much shorter as compared to those produced by sentinel, since foragers were only able to witness the threat in proximity whereas sentinel perceived it from a distance (Sommer 2011). Our results also suggest that the alarm vocalizations produced by foragers, threat call, is monosyllabic with the shortest note duration and functions as a flee call. However, alarm calls like harsh, intermediate alert and alert produced by sentinel or individuals on perch were found to be multisyllabic calls. We speculate that different alarm calls of Jungle Babbler - harsh, intermediate alert and alert are produced in relation to low, intermediate and high urgency of the threat, however, this needs further examination. We speculate that the sentinel soft call could serve the function similar to the ‘watchman song’ reported in other cooperative breeders (Wickler 1985, Manser 1999, Hollén et al. 2008). The ‘watchman song’ is a form of acoustic coordination between vigilant (sentinel) and non-vigilant (foragers) group members, which serves as a proxy for the presence of sentinel on duty for the rest of the group (Wickler 1985). It can be considered as both affiliative and agonistic as it helps in maintaining group cohesion as well as provides information about the predation risk (Kern and Radford 2013).

Overall, our study on the social behaviour and acoustic communication of a sexually monomorphic, cooperatively breeding passerine, Jungle Babbler, extends support to the social complexity hypothesis. The broad distribution of this species in India and its disproportionate investment in vigilance calls make it a good model system to study vigilance behaviour in groups occupying diverse habitats with disparate predation pressures. The quantitative categorization of different call types to note level in our study has rarely been carried out. This along with the information on variation in the acoustic features of notes provides a rich database for future research on understanding rules governing multisyllabic call composition and information encoding in avian vocalizations. While the overall behavioural contexts of the vocalizations are evident from our findings, understanding the proximate function of some of the vocalizations such as the foraging, sentinel soft and intermediate alert will require further studies, including manipulative experiments using field playbacks. This study will be foundational to all such future investigations.

## Acknowledgements

We gratefully acknowledge the help received from Nakul Raj, Ranjit, Gurtej and Sandeep towards conducting the fieldwork and members of BEL, IISER-M for useful discussions. We thank the Director, NIPER for necessary permits to conduct fieldwork at NIPER campus. MJ is grateful to SKM for introducing her to ornithology and to JB for taking over parental care for our fledgling when needed.

## Author contributions

MJ conceived and designed the study. SDY carried out all the field work and acoustic analyses for repertoire size, call characteristics and for behavioural data. SC carried out all the field work and acoustic analyses for note level complexity. SDY, SC and MJ carried out all statistical analyses and wrote the manuscript. All authors approved the final version of the manuscript.

## Data availability

The datasets generated are available as supplementary material

## Compliance with ethical standards

## Ethical approval

Jungle Babblers are listed in Schedule IV under the Indian Wildlife Protection Act (1972) and designated as ‘Least Concern’ by IUCN’s Red List of Threatened Species. This study was conducted with necessary permits (No. 3625) from Department of Forest and Wildlife Preservation, Government of Punjab, India, and with the approval of Institute Animal Ethical Committee (IISER/SAFE/PRT/2018/003), IISER Mohali, India. No animals were harmed or kept in captivity for this study.

