## Supplementary File for "Structural and functional complexity of vocalizations in a cooperatively breeding passerine, Jungle Babbler"

*Corresponding Author

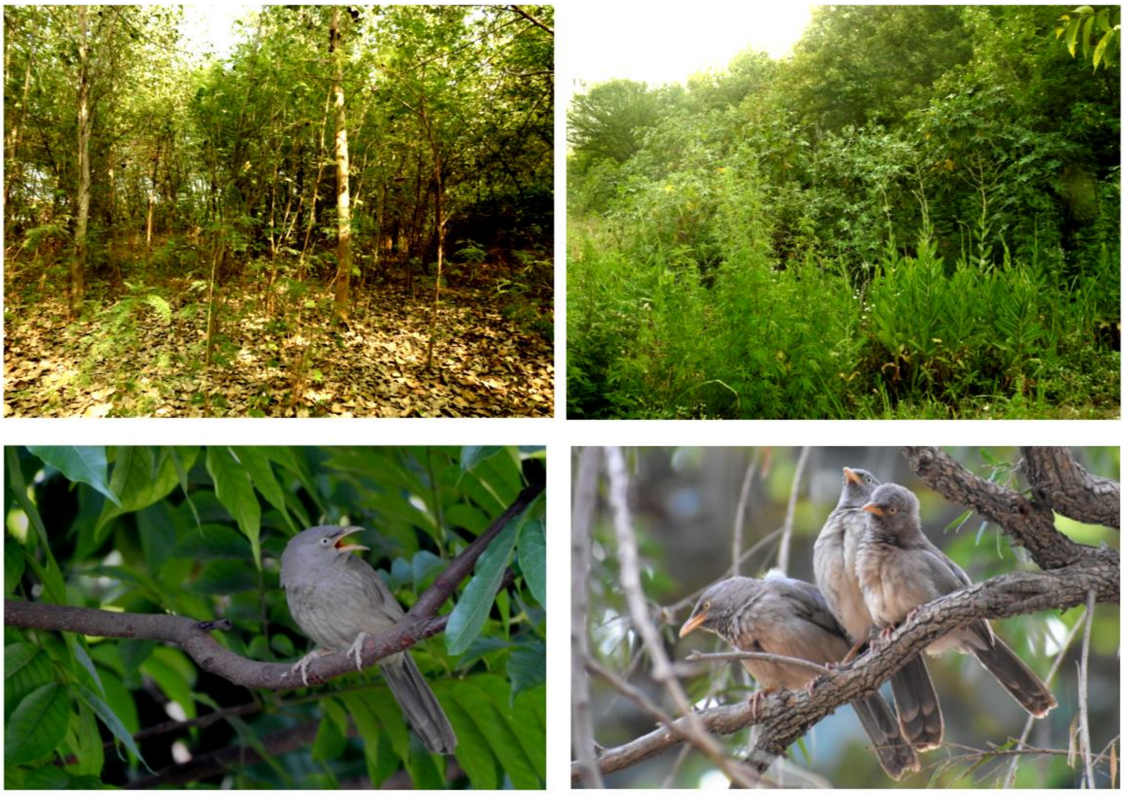

**Fig. S1** Top left and right is JB habitat in the study area. Bottom left: JB perched on *Chukrasia tabularis.* Bottom right: JB perched on a branch of *Callistemon sp.*

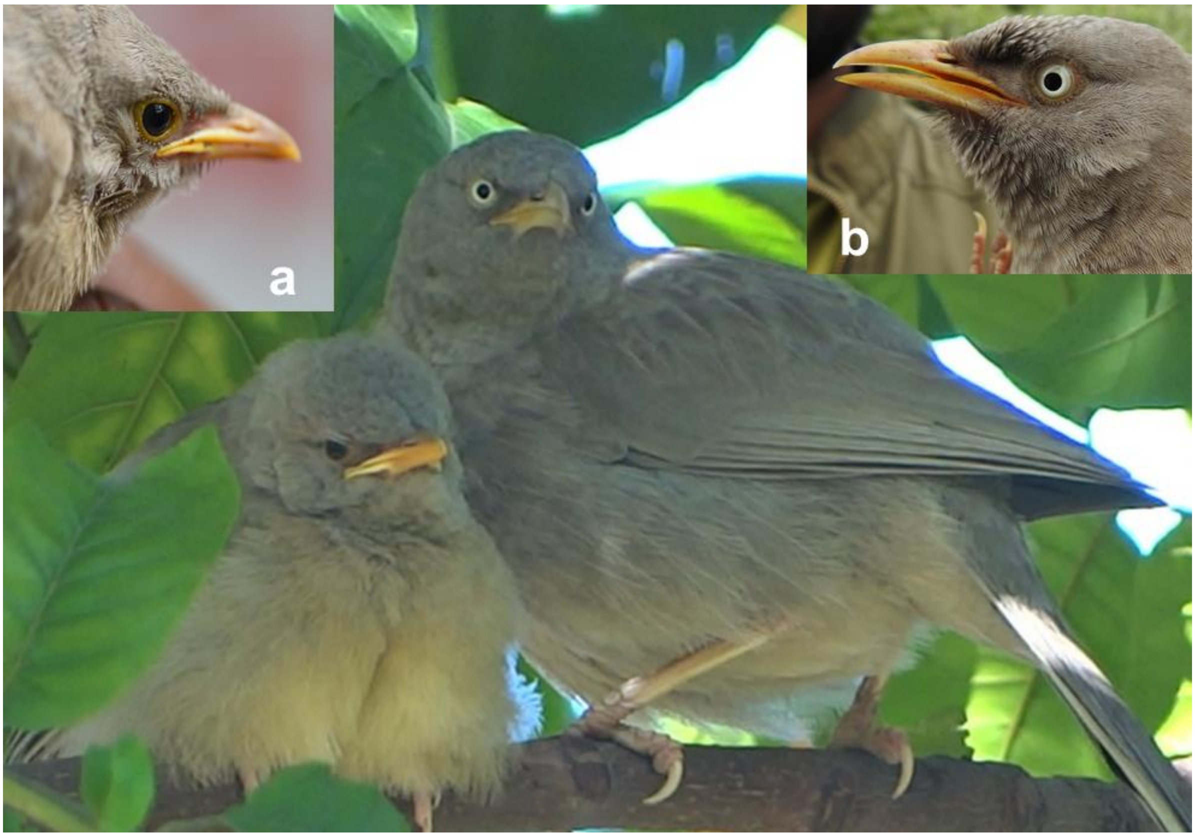


**Fig. S2** Difference in the iris colour of **a** juvenile (black) and **b** adult (pale white) of JB

**Table S1** Statitical analysis of monosyllabic call types of adult of Jungle Babbler. **a** Kruskal-Wallis ANOVA **b** Pairwise comparison using Mann-Whitney U test based on 5 acoustic parameters. 'ns' represents no significant difference (P > 0.05)

**a**

| Call types | Call paramters | $\chi^2$ | df | N | H | p |
|---|---|---|---|---|---|---|
| Monosyllabic | Call duration (s) | 51.852 | 4 | 105 | 85.007 | <0.001 |
| | Frequency 5% (Hz) | 53.845 | 4 | 105 | 73.024 | <0.001 |
| | Frequency 95% (Hz) | 51.832 | 4 | 105 | 56.992 | <0.001 |
| | Bandwidth 90% (Hz) | 50.488 | 4 | 105 | 60.580 | <0.001 |
| | Peak frequency (Hz) | 21.184 | 4 | 105 | 42.785 | <0.001 |

**b**

| Call type | Frequency 5% (Hz) | Frequency 95% (Hz) | Bandwidth 90% (Hz) | Peak frequency (Hz) | Call duration (s) |
|---|---|---|---|---|---|
| Contact vs Sentinel soft | <0.001 | <0.001 | <0.001 | <0.01 | <0.01 |
| Contact vs Threat | <0.001 | <0.001 | <0.001 | <0.01 | <0.001 |
| Contact vs Distress | ns | <0.001 | <0.001 | ns | <0.001 |
| Contact vs Foraging | <0.001 | <0.001 | <0.001 | <0.001 | <0.001 |
| Sentinel soft vs Threat | <0.001 | <0.01 | ns | <0.001 | <0.05 |
| Sentinel soft vs Distress | <0.001 | ns | ns | <0.01 | <0.001 |
| Sentinel soft vs Foraging | <0.05 | <0.01 | <0.01 | <0.01 | <0.001 |
| Threat vs Distress | <0.001 | ns | ns | <0.05 | <0.001 |
| Threat vs Foraging | <0.001 | ns | <0.001 | <0.001 | <0.001 |
| Distress vs Foraging | <0.001 | ns | <0.001 | <0.001 | <0.001 |

**Table S2** A summary of pairwise comparison of monosyllabic call types of juvenile using t-test based on 5 different acoustic parameters. 'ns' represents no significant difference (P > 0.05)

| Call types | Call parameters | t | df | p |
|---|---|---|---|---|
| Juvenile | Call duration (s) | 5.595 | 22 | <0.001 |
| | Frequency 5% (Hz) | -1.066 | 22 | ns |
| | Frequency 95% (Hz) | 1.464 | 22 | ns |
| | Bandwidth 90% (Hz) | 1.701 | 22 | ns |
| | Peak frequency (Hz) | 0.988 | 22 | ns |

**Table S3** Statistical analysis of multisyllabic call types. **a** Analysis of Variance **b** Kruskal-Wallis ANOVA. Pairwise comparison using **c** t-test and **d** Man-Whitney U test based on 7 different acoustic parameters. 'ns' represents no significant difference (P > 0.05)

**a**

| **Call types** | **Call parameters** | **F** | **df** | **p** |
|---|---|---|---|---|
| Multisyllabic | Bandwidth 90 (Hz) | 10.336 | 5 | <0.001 |

**b**

| **Call types** | **Call paramters** | $\chi^2$ | **df** | **N** | **H** | **p** |
|---|---|---|---|---|---|---|
| Multisyllabic | Number of notes | 69.904 | 5 | 152 | 91.538 | <0.001 |
| | Inter-note interval (s) | 41.304 | 5 | 137 | 55.963 | <0.001 |
| | Note duration (s) | 52.315 | 5 | 152 | 56.548 | <0.001 |
| | Frequency 5% (Hz) | 89.936 | 5 | 152 | 95.239 | <0.001 |
| | Frequency 95% (Hz) | 58.862 | 5 | 152 | 67.894 | <0.001 |
| | Peak frequency (Hz) | 74.410 | 5 | 152 | 90.520 | <0.001 |

**c**

| **Call types** | | **Call parameters** | **t** | **df** | **p** |
|---|---|---|---|---|---|
| Multisyllabic | Alert vs Harsh | Bandwidth 90% (Hz) | 3.310 | 59 | <0.01 |
| | Alert vs Intermediate alert | | 3.733 | 40 | <0.001 |
| | Alert vs Flight | | 7.555 | 52 | <0.001 |
| | Alert vs Prompt flight | | 4.742 | 43 | <0.001 |
| | Alert vs Prompt | | 0.287 | 40 | ns |
| | Harsh vs Intermediate alert | | -0.014 | 55 | ns |
| | Harsh vs Flight | | 3.462 | 67 | <0.001 |
| | Harsh vs Prompt flight | | 1.673 | 58 | ns |
| | Harsh vs Prompt | | -2.183 | 55 | <0.05 |
| | Intermediate alert vs Flight | | 3.534 | 48 | <0.001 |
| | Intermediate alert vs Prompt flight | | 1.649 | 39 | ns |
| | Intermediate alert vs Prompt | | -1.993 | 36 | ns |
| | Flight vs Prompt flight | | -1.227 | 51 | ns |
| | Flight vs Prompt | | -4.833 | 48 | <0.001 |
| | Prompt flight vs Prompt | | -3.070 | 39 | <0.01 |

**d**

| Call type | Frequency 5% (Hz) | Frequency 95% (Hz) | Peak frequency (Hz) | No. notes in a call (s) | Inter-note interval (s) | Call duration (s) |
|---|---|---|---|---|---|---|
| Alert vs Harsh | <0.05 | <0.001 | <0.01 | <0.001 | <0.01 | <0.01 |
| Alert vs Intermediate alert | ns | <0.01 | ns | ns | <0.01 | <0.01 |
| Alert vs Flight | <0.001 | <0.001 | <0.001 | ns | ns | <0.01 |
| Alert vs Prompt flight | <0.001 | <0.001 | <0.001 | <0.001 | <0.05 | ns |
| Alert vs Prompt | <0.001 | ns | <0.001 | ns | <0.05 | <0.05 |
| Harsh vs Intermediate alert | <0.001 | ns | <0.05 | <0.001 | ns | <0.001 |
| Harsh vs Flight | <0.001 | <0.001 | <0.001 | <0.001 | <0.001 | ns |
| Harsh vs Prompt flight | <0.001 | <0.001 | <0.001 | <0.001 | <0.001 | <0.01 |
| Harsh vs Prompt | <0.001 | ns | <0.001 | <0.001 | <0.001 | ns |
| Intermediate alert vs Flight | <0.001 | <0.001 | <0.001 | <0.05 | <0.001 | <0.001 |
| Intermediate alert vs Prompt flight | <0.001 | <0.001 | <0.001 | <0.001 | <0.001 | <0.001 |
| Intermediate alert vs Prompt | <0.001 | ns | <0.001 | <0.01 | <0.001 | <0.001 |
| Flight vs Prompt flight | ns | ns | ns | <0.001 | ns | <0.001 |
| Flight vs Prompt | ns | <0.001 | <0.05 | <0.05 | <0.01 | ns |
| Prompt flight vs Prompt | ns | <0.01 | ns | <0.01 | ns | <0.001 |

**Table S4** A summary of pairwise comparison of chorus call types using **a** t-test and **b** Mann-Whitney U test based on 4 different acoustic parameters. 'ns' represent no significant difference (P > 0.05)

**a**

| Call types | Call parameters | t | df | p |
|---|---|---|---|---|
| Chorus | Frequency 95% (Hz) | 1.513 | 20 | ns |
| | Bandwidth 90% (Hz) | 0.555 | 20 | ns |
| | Peak frequency (Hz) | 0.94 | 20 | ns |

**b**

| Call types | Call parameters | p |
|---|---|---|
| Chorus | Frequency 5% (Hz) | ns |

**Table S5** Statistical analysis of note types within a multisyllabic call. **a** Analysis of Variance **b** Kruskal-Wallis ANOVA and **c** pairwise comparison using Mann-Whitney U test based on 5 different acoustic parameters. 'ns' represent no significant difference (P > 0.05)

**a**

| Note types | Call parameters | F | df | p |
|---|---|---|---|---|
| Prompt P1;P2;P3;P4 | Frequency 5% (Hz) | 1.741 | 3 | ns |
| Flight F1;F2;F3 | Frequency 5% (Hz) | 0.957 | 2 | ns |

**b**

| Note types | Call Parameters | $\chi^2$ | df | N | H | p |
|---|---|---|---|---|---|---|
| Prompt P1;P2;P3;P4 | Note duration (s) | 15.345 | 3 | 80 | 23.484 | <0.0001 |
| | Frequency 95% (Hz) | 1.350 | 3 | 80 | 1.634 | ns |
| | Bandwidth 90% (Hz) | 1.200 | 3 | 80 | 2.100 | ns |
| | Peak frequency (Hz) | 0.550 | 3 | 80 | 1.523 | ns |
| Flight F1;F2;F3 | Note duration (s) | 41.202 | 2 | 150 | 62.736 | <0.0001 |
| | Frequency 95% (Hz) | 29.498 | 2 | 150 | 25.882 | <0.0001 |
| | Bandwidth 90% (Hz) | 17.920 | 2 | 150 | 16.080 | <0.0001 |
| | Peak frequency (Hz) | 0.377 | 2 | 150 | 2.410 | ns |
| Flight prompt PF1;PF2;PF3;PF4; PF5;PF6;PF7 | Note duration (s) | 33.944 | 6 | 354 | 57.022 | <0.0001 |
| | Frequency 5% (Hz) | 3.202 | 6 | 354 | 10.902 | ns |
| | Frequency 95% | 10.751 | 6 | 354 | 6.457 | ns |
| | Bandwidth 90% | 3.756 | 6 | 354 | 2.848 | ns |
| | Peak frequency (Hz) | 15.960 | 6 | 354 | 14.186 | ns |
| Alert A1;A2;A3 | Note duration (s) | 133.498 | 3 | 155 | 126.335 | <0.0001 |
| | Frequency 5% (Hz) | 87.829 | 3 | 155 | 103.532 | <0.0001 |
| | Frequency 95% | 4.955 | 3 | 155 | 3.998 | ns |
| | Bandwidth 90% | 24.443 | 3 | 155 | 36.405 | <0.0001 |
| | Peak frequency (Hz) | 2.146 | 3 | 155 | 2.907 | ns |
| Harsh H1 ;H2;H3;H4 | Note duration (s) | 326.854 | 3 | 705 | 504.085 | <0.0001 |
| | Frequency 5% (Hz) | 1.445 | 3 | 705 | 1.678 | ns |
| | Frequency 95% | 4.613 | 3 | 705 | 8.279 | <0.05 |
| | Bandwidth 90% | 11.506 | 3 | 705 | 13.677 | <0.01 |
| | Peak frequency (Hz) | 16.146 | 3 | 705 | 13.326 | <0.01 |
| Intermediate alert M1;M2;M3;M4;M 5;M6 | Note duration (s) | 125.431 | 5 | 464 | 200.421 | <0.0001 |
| | Frequency 5% (Hz) | 166.164 | 5 | 464 | 197.753 | <0.0001 |
| | Frequency 95% | 27.338 | 5 | 464 | 33.055 | <0.0001 |
| | Bandwidth 90% | 58.473 | 5 | 464 | 70.553 | <0.0001 |
| | Peak frequency (Hz) | 57.580 | 5 | 464 | 62.028 | <0.0001 |

**c**

| Parameters Note types | Frequency 5% (Hz) | Frequency 95% (Hz) | Bandwidth 90% (Hz) | Peak frequency (Hz) | Note duration (s) |
|---|---|---|---|---|---|
| P1 vs P2 | ns | ns | ns | ns | <0.05 |
| P1 vs P3 | ns | ns | ns | ns | <0.01 |
| P1 vs P4 | ns | ns | ns | ns | <0.001 |
| P2 vs P3 | ns | ns | ns | ns | ns |
| P2 vs P4 | ns | ns | ns | ns | <0.01 |
| P3 vs P4 | ns | ns | ns | ns | <0.05 |
| F1 vs F2 | ns | <0.001 | <0.01 | ns | <0.001 |
| F1 vs F3 | ns | <0.001 | <0.001 | ns | <0.001 |
| F2 vs F3 | ns | ns | ns | ns | <0.001 |
| PF1 vs PF2 | ns | ns | ns | ns | <0.01 |
| PF1 vs PF3 | ns | ns | ns | ns | <0.001 |
| PF1 vs PF4 | ns | ns | ns | <0.05 | <0.001 |
| PF1 vs PF5 | ns | ns | ns | ns | <0.001 |
| PF1 vs PF6 | ns | ns | ns | <0.01 | <0.001 |
| PF1 vs PF7 | ns | ns | ns | <0.05 | <0.001 |
| PF2 vs PF3 | ns | ns | ns | ns | ns |
| PF2 vs PF4 | ns | ns | ns | ns | <0.01 |
| PF2 vs PF5 | ns | ns | ns | ns | <0.01 |
| PF2 vs PF6 | ns | ns | ns | <0.05 | <0.001 |
| PF2 vs PF7 | ns | ns | ns | ns | <0.01 |
| PF3 vs PF4 | ns | ns | ns | ns | ns |
| PF3 vs PF5 | ns | ns | ns | ns | ns |
| PF3 vs PF6 | ns | ns | ns | <0.05 | <0.001 |
| PF3 vs PF7 | ns | ns | ns | ns | ns |
| PF4 vs PF5 | ns | ns | ns | ns | ns |
| PF4 vs PF6 | ns | ns | ns | ns | <0.01 |
| PF4 vs PF7 | ns | ns | ns | ns | ns |
| PF5 vs PF6 | ns | ns | ns | ns | <0.05 |
| PF5 vs PF7 | ns | ns | ns | ns | ns |
| PF6 vs PF7 | ns | ns | ns | ns | ns |
| H1 vs H2 | ns | ns | ns | ns | <0.001 |
| H1 vs H3 | ns | ns | <0.05 | <0.05 | <0.001 |
| H1 vs H4 | ns | ns | ns | <0.001 | <0.001 |

| | | | | | |
|---|---|---|---|---|---|
| H2 vs H3 | ns | <0.01 | <0.001 | ns | <0.001 |
| H2 vs H4 | ns | ns | ns | <0.01 | <0.001 |
| H3 vs H4 | ns | ns | ns | <0.01 | <0.001 |
| A1 vs A2 | <0.001 | ns | ns | ns | <0.001 |
| A1 vs A3 | <0.001 | ns | <0.001 | ns | <0.001 |
| A1 vs A4 | <0.001 | ns | <0.001 | ns | <0.001 |
| A2 vs A3 | <0.001 | ns | <0.01 | ns | <0.001 |
| A2 vs A4 | <0.001 | ns | <0.01 | ns | <0.001 |
| A3 vs A4 | <0.001 | ns | ns | ns | <0.001 |
| M1 vs M2 | <0.05 | ns | ns | ns | <0.001 |
| M1 vs M3 | <0.01 | <0.01 | <0.001 | ns | <0.001 |
| M1vs M4 | <0.001 | ns | <0.001 | <0.01 | <0.001 |
| M1 vs M5 | <0.05 | ns | <0.05 | ns | ns |
| M1 vs M6 | <0.001 | ns | <0.05 | <0.001 | <0.001 |
| M2 vs M3 | ns | <0.001 | <0.001 | ns | <0.001 |
| M2 vs M4 | <0.001 | ns | <0.001 | <0.001 | <0.001 |
| M2 vs M5 | ns | ns | ns | ns | <0.001 |
| M2 vs M6 | <0.001 | ns | <0.05 | <0.001 | ns |
| M3 vs M4 | <0.001 | <0.001 | ns | <0.01 | <0.001 |
| M3 vs M5 | ns | ns | ns | <0.05 | <0.001 |
| M3 vs M6 | <0.001 | <0.001 | <0.001 | <0.001 | <0.001 |
| M4 vs M5 | <0.001 | ns | <0.01 | ns | <0.001 |
| M4 vs M6 | <0.001 | <0.01 | <0.001 | ns | <0.001 |
| M5 vs M6 | <0.01 | ns | ns | ns | <0.01 |
